# Comparative genome analysis of 12 *Shigella sonnei* strains: virulence, resistance, and their interactions

**DOI:** 10.1101/544387

**Authors:** Zuobin Zhu, Liang Wang, Feng Gu, Ying Li, Heng Zhang, Ying Chen, Jiajia Shi, Ping Ma, Bing Gu

## Abstract

Shigellosis is a highly infectious disease that are mainly transmitted via faecal-oral contact of the bacteria *Shigella.* Four species have been identified in *Shigella* genus, among which *S. flexneri* is used to be the most prevalent species globally and commonly isolated from developing countries. However, it is being replaced by *S. sonnei* that is currently the main causative agent for dysentery pandemic in many emerging industrialized countries such as Asia and the Middle East with unclear reasons. For a better understanding of *S. sonnei* virulence and antibiotic resistance, we sequenced 12 clinical *S. sonnei* strains with varied antibiotic-resistance profiles collected from four cities in Jiangsu Province, China. Phylogenomic analysis clustered antibiotic sensitive and resistant *S. sonnei* into two distinct groups while pangenome analysis reveals the presence and absence of featured genes in each group. Screening of 31 classes of virulence factors found out that type 2 secretion system is doubled in resistant strains. Further principle component analysis based on the interactions between virulence and resistance indicated that abundant virulence factors are associated with higher resistant phenotypes. The result present here is based on statistical analysis of a small sample size and serves basically as a guidance for further experimental and theoretical studies.

## Introduction

Shigellosis is a life-threatening diarrheal infection (dysentery) and is currently a major global burden. Although four species fall into the genus of *Shigella, S. sonnei* and *S. flexneri* are now accounting for the most dysentery worldwide while the other two strains *S. dysenteriae* and *S. boydii* experience inexplicable absence^1^. Although *S. flexneri* is the leading cause of endemic shigellosis in developing countries with around 75% episodes, *S. sonnei* is increasing in many rapidly developing countries due to the improvement of socioeconomic conditions^1,2^. The rapid expansion of *S. sonnei* have been attributed to a couple of speculative reasons, such as passive immunization caused by *Plesiomonas shigelloides* and protected niche provided by ubiquitous amoeba species *Acanthamoeba castellaniii^2^.* In addition, *S. sonnei* is more capable of incorporating functional virulence factors (VFs) and/or adept at acquiring antibiotic resistance (AR) from other bacteria^1,3^. Hence, enhanced survival advantages. However, a recent study showed that *S. sonnei* is not able to use amoebae as a protective host to enhance its environmental survival, which makes the explanation of *A. castellaniii* protection for *S. sonnei* expansion compromised^4^. On the other hand, antibiotic resistance is thought to be associated with a fitness cost^5^. Thus, resistance and virulence are historically thought to be negatively correlated^6^. However, recent studies indicated that antibiotic selection pressure and genetic associations could lead to the co-occurrence of resistance and virulence in multiple pathogenic bacteria^7^.

In this study, we sequenced 12 clinically isolated strains with diverse antibiotic resistance phenotypes. Sequenced *S. sonnei* genomes were aligned and compared with the reference strain *S. sonnei* 53G. The relationship between the antibiotic resistance and virulence were studied by combining antibiotic resistance profiles with the distribution of putative virulence factors. Here, what we mean by virulence factors is gene products that enable a microorganism to establish itself on or within a host of a particular species, facilitating its abilities to cause diseases, which are divided into 4 categories and 31 functional groups, such as bacterial toxins, cell surface proteins, and hydrolytic enzymes, *etc^8^.* All the virulence factors come from 32 major bacterial pathogens including the *Shigella* genus. By screening and comparing the genomes of sensitive and antibiotic resistant *Shigella sonnies* using this hierarchical set of VF sequence models, we attempted to quantify virulence by the number of virulence factors in specific functional groups. Principal component analysis was then performed to cluster the resistant and sensitive strains, separately, via incorporating both the number of virulence factors and the degree of antibiotic resistance. Genes uniquely associated with sensitive and resistant strains were also studied. Based on these analyses, we could get a better understanding of how virulence and antibiotic resistance are interacted.

## Results

### Genome annotation and comparison

General features of 12 *Shigella sonnei* genomes are presented in Supplementary Table 1, which include information about resistance profiles to 9 previously mentioned antibiotics, genome size, contigs, coding sequences, and RNAs (tmRNA, tRNA, rRNA). All strains were sensitive to Norfloxacin and most strains were susceptible to Amoxicillin/Clavulanic acid. Genome size ranges from 4.49 Mbps to 4.76Mbps. The number of CDSs ranges from 4328 (S13029) to 4646 (S14049). All strains have a single tmRNA coding gene. The number of ribosome RNA (rRNA) and transfer RNA (tRNA) among strains varies slightly with no significant difference. All the genomes were aligned against reference genome S. sonnei G53 in circular form via BRIG in terms of distribution of GC content and sequence similarity in Supplementary Figure 1^9^. Absence of large blocks in both sensitive and resistant strains were observed, which requires further investigation for their biological meanings.

### Pan- and phylo-genomic analysis

The total pan-genome for the 12 *S. sonnei* strains include 5608 protein CDSs. Of those, 3893 (69.42% of total CDSs) are core genes across all 12 species while 1715 (30.58% of total CDSs) constitute the accessory fractions, which are unique to each genome. Strain S13029 has the lowest number of the unique genes (484 CDSs) and S14031 has the highest number of unique genes (803 CDSs) (Supplementary Figure 2). Interestingly, both strains are completely sensitive to or only resistant to one of all tested antibiotics. Further comparison of all the 12 *S. sonnei* strains give complete map of gene presence and absence in each genome (Dataset 1). By comparing sensitive strains with resistant strains, unique genes associated with the two groups were identified and annotated based on sequence homology (**Supplementary Table 2**), respectively. These genes could serve as a guidance for a better understanding of the differences between the two *S. sonnei* groups in terms of their virulence and resistance. Phylogenomic analysis based on the concatenation of 3893 core genes clearly classified the 12 strains into two distinct groups (Figure 1). Antibiotic sensitive strains S13029 and S14031 are distantly related with other 10 antibiotic resistant strains, which suggested distinct difference between the two groups in terms of evolutionary pathway.

**Figure 1.**
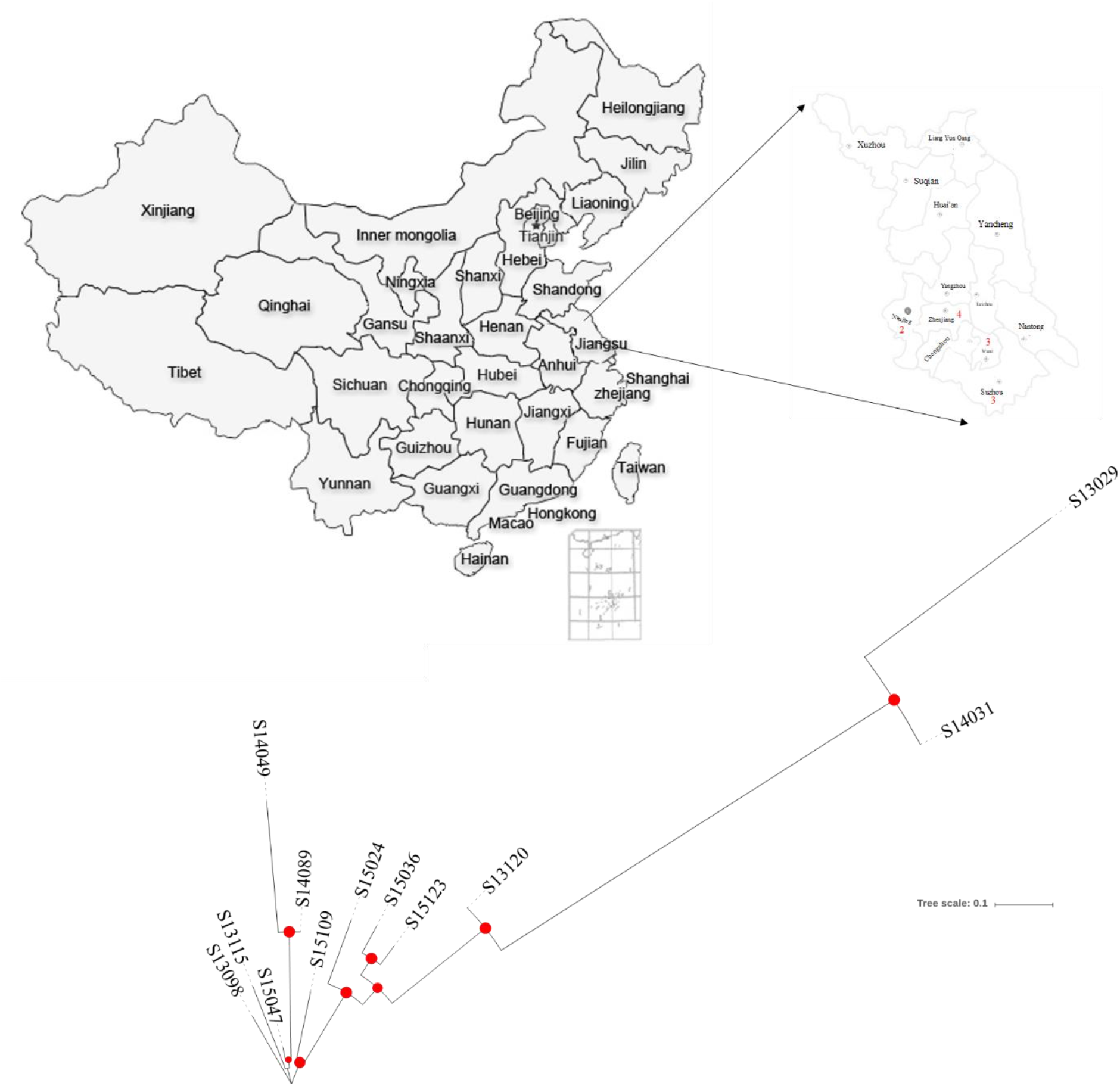
Phylogenomic analysis of 12 clinically isolated *Shigella sonnei* strains with different antibiotic resistant profiles based on 3893 core genes via FastTree. S13029 is completely sensitive to all 9 tested antibiotics, that is, Amoxicillin/Clavulanic acid (AMC), Ceftiophene (CFT), Cefotaxime (CTX), Gentamicin (GEN), Nalidixic acid (NAL), Norfloxacin (NOR), Tetracycline (TBT), and compound Sulfamethoxazole (SMZ) while S14031 is only resistant to SMZ. The other ten strains are resistant to at least five out of the nine antibiotics. All the strains were isolated from 4 municipal cities (Nanjing, Suzhou, Wuxi, Zhenjiang) in Jiangsu Province, China, the geographic distribution of which was depicted in the figure with red numbers. Apparent clusters could be observed for sensitive and resistant strains, respectively. Branch length represents evolutionary distance among strains. Filled red dots indicate bootstrapping value greater than 90%.

### Interactions between virulence and resistance

In order to investigate the correlations between resistance and virulence, we divided all virulence factors into 4 categories and 31 groups based on VFDB’s instructions as specified above^8^, which were then used to identify the presence and absence of virulence factors in all studies genomes. Distribution patterns of virulence factors and their abundance in each strain were presented in Table 1. Nine groups of virulence factors are completely missing in all *S. sonnei* while three groups of virulence factors are equally distributed in each genome. Among the rest 19 groups of virulence factors, Type 4 pili and T2SS shows apparent differences between sensitive and resistant groups. If only strain S13029 was considered, the number of most groups of virulence factors is reduced except for the three groups Chaperone usher, Flagella, and T3SS. Principal component analysis was also performed by combining antibiotic resistance profiles and distributions of virulence factors, according to which, sensitive strains S13029 and S14031 are separately clustered when compared with other resistant strains (Supplementary Figure 3). In addition, S13029 is most isolated due to its tight correlation between reduced virulence and resistance loss.

**Table 1.**
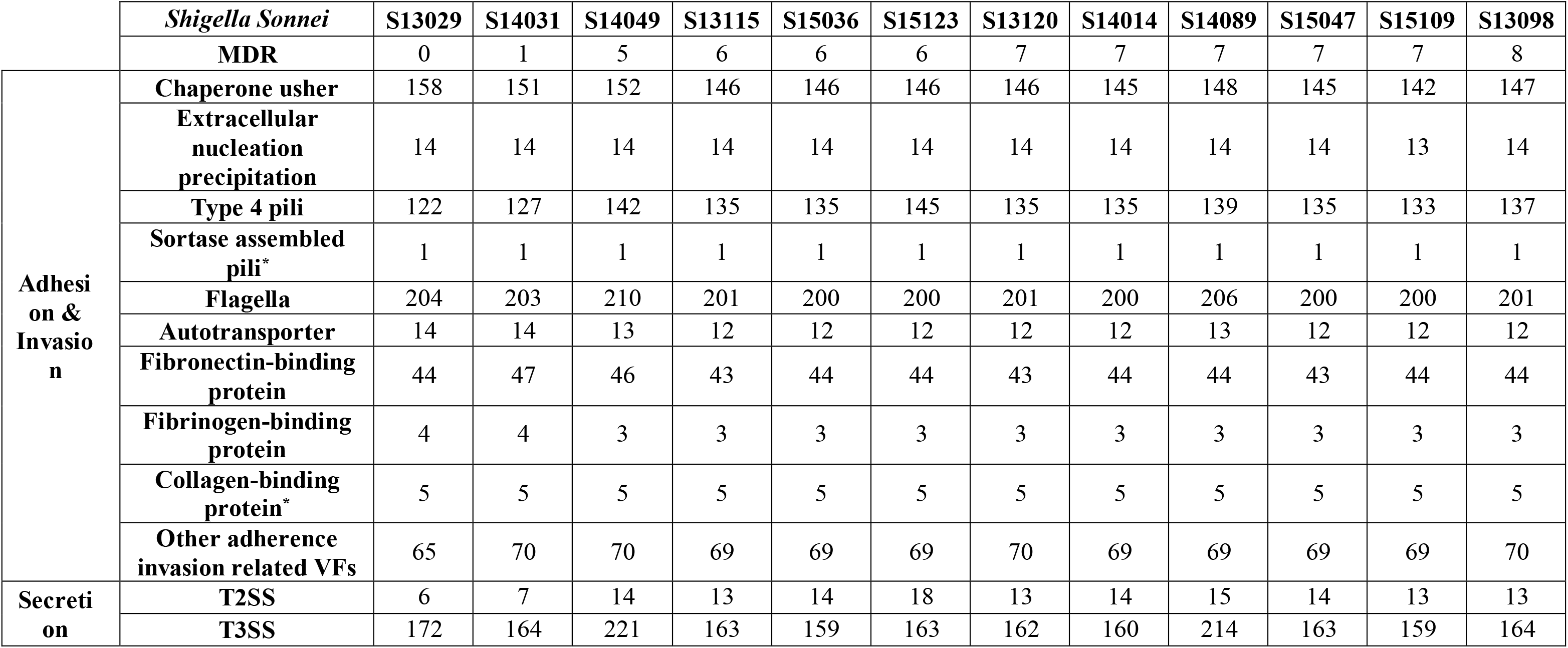
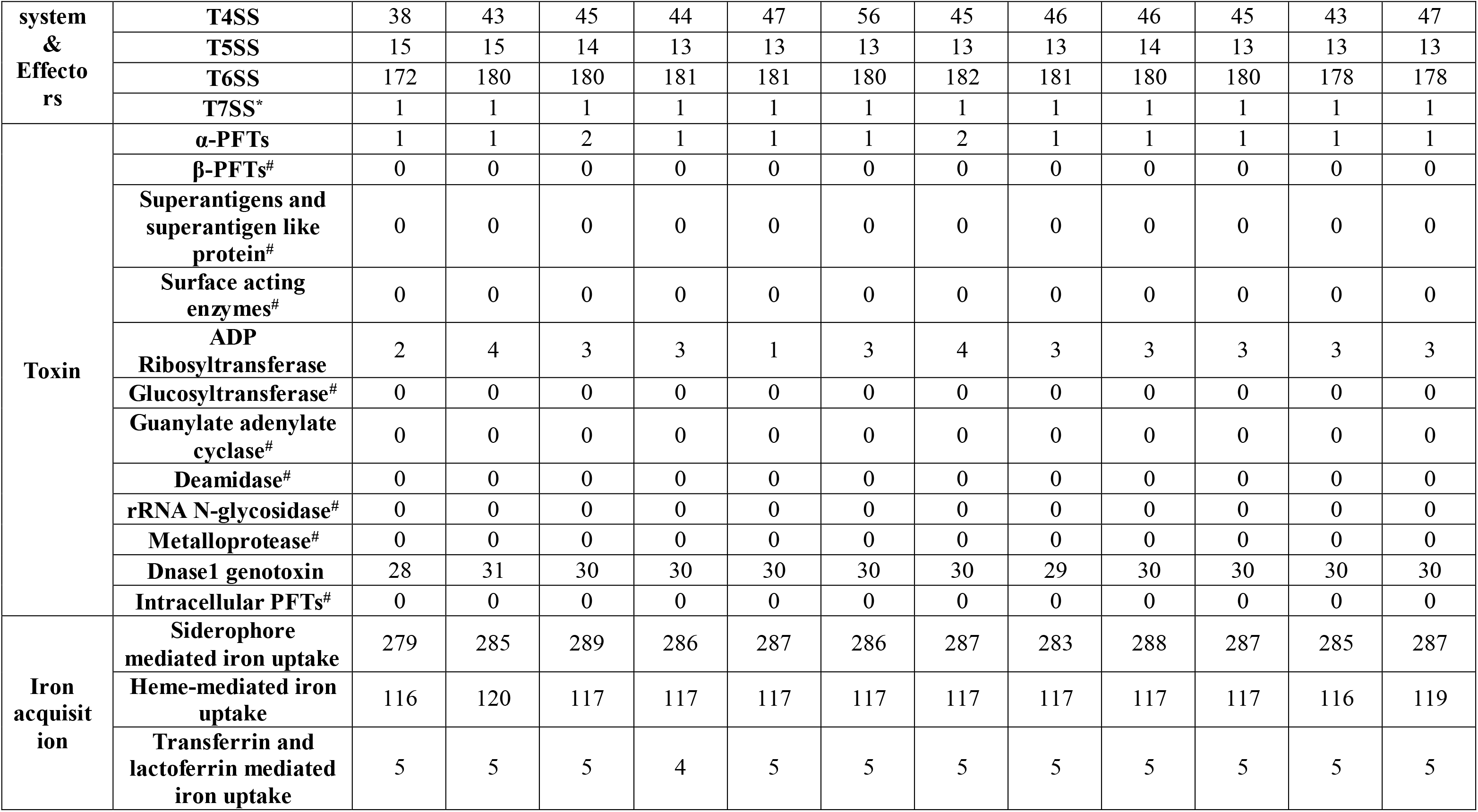
Distribution patterns of 4 categories of virulence factors that belong to 31 groups among 12 *Shigella sonnei* strains in terms of antibiotic resistance. Sensitive strains have no resistance or only resist to one antibiotics. The four categories of VFs are Adhesion & Invasion, Secretion system & Effectors, Toxin, Iron acquisition. MDR strains have more than one resistance. PFT: Pore-forming toxins. #Nine groups of virulence factors are not present in all *S. sonnei* strains. *Three groups of virulence factors have equal number of virulence factors in all strains.

Further analysis in terms of unique genes associated with sensitive strains or resistant strains were also analysed, which identified 26 genes specifically associated *with Shigella sonnei* sensitive strains and 39 unique genes with resistant strains (**Supplementary Table 2**). Investigation into these genes could shed light on a better understanding of physiological differences between resistant and sensitive *S. sonnei* strains. Genes with no assigned names are generally hypothetical proteins and will not be considered in this study.

## Discussion

In this study, 12 newly isolated *S. sonnei* strains were sequenced and assembled into complete genomes, which were then well annotated and thoroughly analysed. It is generally accepted that antibiotic resistance causes fitness costs such as slow growth rate^10^ and virulence attenuation ^11^. However, recent studies are challenging this view by providing evidence in which drug resistance leads to increased pathogenicity^11^. This study performed a bioinformatics analysis by focusing on the interplays between virulence and resistance based on 12 newly sequenced *S. sonnei* strains. Through the distribution of functional groups of virulence factors in *S. sonnei* strains, specific patterns were observed. Nine virulence factor groups were completely absent in all S. sonnei strains while another three groups have equal number of virulence factors in all strains (Table 1). Thus, their interactions with antibiotic resistance were not considered. As for the rest 19 groups of virulence factors, type 4 pili and T2SS genes are skewedly present in more resistant *S. sonnei* strains when compared with the two sensitive strains S13029 and S14031. Type 4 pili is a bacterial extracellular appendage essential for attachment to host cells while T2SS is responsible for the secretion of numerous degradative enzymes and toxins for bacterial survival^12^. The reasons for their association with high antibiotic resistance are worthy of further exploration. Considering the versatility of virulence mechanisms, PCA was performed to study the interactions between virulence and resistance. It was clearly shown that completely sensitive strain S13029 and single-resistant strain S14031 are distantly separated from other resistant strains that cluster together (Supplementary Figure 3). The statistical analysis provided a theoretical support for the view that high virulence is associated with high resistance^11^. However, what the mechanisms are behind this putative relationship is still unclear and requires more efforts to solve the puzzle.

On the other hand, pangenome analysis also revealed vast genome heterogenicity within the same strains of *S. sonnei* according to the identified core and cloud genes (Supplementary Figure 1). These cloud genes, more precisely known as strain specific genes, could reveal bacterial characteristics and dynamics, leading to a better understanding of physiological, pathological, and epidemiological features of *S. sonnei^13^.* In specificity, all unique genes that are respectively associated with sensitive and resistant strains were listed in **Supplementary Table 2**. Apparent differences could be observed from the comparisons of these gene functions. However, more theoretical and experimental studies should be performed on a larger scale of datasets in order to get a better understanding of how virulence and resistance are interacted.

## Conclusion

In this study, *S. sonnei* sensitive strains was clearly separated from resistant strains based on phylogenomic analysis. By combining antibiotic resistance profiles with the distribution of putative virulence factors, we explored the relationships between virulence and resistance.

Reduced number of putative virulence factors, especially for the group of Type 4 pili and T2SS, was observed to be related with antibiotic sensitivity. Principal component analysis also clustered the resistant and sensitive strains, separately, via incorporating both the abundance of virulence factors and the degree of antibiotic resistance. Finally, genes uniquely associated with sensitive and resistant strains were also reported. In order to better understand how resistance and virulence are interacted, further theoretical and experimental studies should be performed.

## Methods and materials

### Bacterial isolates and DNA extraction

12 *Shigella sonnei* strains were isolated from different patients with either diarrhea or dysentery in four cities in Jiangsu province, China. Resistance profile to 9 antibiotics (AMC, CFT, CTX, GEN, NAL, NOR, TBT and SMZ) as previously described for each strain was provided by Jiangsu Provincial CDC based on routine screening. Isolates of *Shigella flexneri* were plated on trypticase soya agar (TSA). Picked-up single colony was then inoculated in 5ml trypticase soya broth (TSB) and incubated overnight at 37°C with shaking rate of 200 rpm. DNA isolation was performed using Easy-DNA^TM^ Kit for genomic DNA isolation (Invitrogen Life Technologies, Carlsbad, CA, USA)

### Genome sequencing, assembly, and annotation

Sequencing and assembly. Genomes of the 12 Shigella sonnei strains were performed using Illumina Hiseq4000 by generating multiplexed paired-end libraries with an average insert size of 300 bp. In order to obtain more accurate and reliable results in subsequent bioinformatics analysis, the raw data will be treated: (1) Read1 selects 1 bp-150 bp, read2 selects 1 bp-150 bp; (2) Remove reads with a certain proportion of low quality (20) bases (40% as default, parameter setting at 60 bp); (3) Remove reads with a certain proportion of Ns (10% as default, parameter setting at 15 bp); (4) Remove adapter contamination (15 bp overlap between adapter and reads as default, parameter setting at 15 bp); (5) Remove duplication contamination. We assemble the short reads into genome sequence using SOAPdenovo^14^. De novo assembly of human genomes with massively parallel short read sequencing^15^. Key parameter K setting at 89 is determined by optimal assembly result for all the 12 strains. Then the assembly result is local assembled and optimized according to paired-end and overlap relationship via mapping reads to Contig. The detailed description of assembly results were provided in Supplementary Table 4.

Obtained sequences were assessed via FastQC, assembled via SPAdes^16^, and reordered via MAUVE^17^ based on reference genome *S. sonnei* 53G by following Edwards and Holt’s beginner’s guide to comparative bacterial genome analysis using next-generation sequence data (Version 2)^18^. For the annotation process, assembled DNA sequences of the drafted genomes from the 12 isolates were run through an automatic annotation pipeline via Prokka (rapid prokaryotic genome annotation)^19^.

### Pan- and phylo-genomic analysis

Core-/pan-genome analysis was performed by using standalone software Roary^18^. Core genes (99≤strains≤100%), soft core genes (95≤strains<99%), shell genes (15≤strains<95%) and cloud genes (0≤strains<15%) were calculated. Core and unique genes in the genomes were illustrated in Venn diagram. Presence and absence of all genes in each genome were summarized in Supplementary Table 1. *S. sonnei* genomes were visualized in circular form by comparing to the reference genome *S. sonnei* 53G via standalone software BRIG^9^. A Newick tree for 12 *Shigella sonnei* strains, was generated based on 3893 core genes in each genome by the phylogenomic analysis package FastTree^21^. The tree was then visualized through online webserver interactive Tree of Life (iTOL)^22^.

### Interactions between virulence and resistance

31 functional groups of bacterial virulence factors belonging to four categories were downloaded from the Virulence Factor Database (VFDB)^8^ and used to screen translated CDSs of the 12 *Shigella sonnei* strains via phmmer command (full-length alignment with e-value less than 1e-5) in HMMER package^23^. For each group in each proteome, multiple homologous sequences of virulence factors were found, which were then processed to get rid of redundant sequences. MDR (resistance to more than 1 antibiotics) and sensitive *S. sonnei* strains (resistance to 0 or 1 antibiotics) were compared in terms of the abundance of specific groups of virulence factors via Python scripts (available under request). Principal component analysis was performed by incorporating both antibiotic resistance profiles and distribution of virulence factors.

## Acknowledgements

This work was supported by the National Natural Science Foundation of China (81871734, 81471994), Jiangsu Provincial Natural Science Foundation (BK20151154, BK20180997), Jiangsu Provincial Medical Talent (ZDRCA2016053), Six Talent Peaks Project of Jiangsu Province (WSN-135), Advanced Health Talent of Six-one Project of Jiangsu Province (LGY2016042), and Jiangsu Provincial Commission of Health and Family Planning Research Project (H201631), Startup Foundation for Excellent Researchers at Xuzhou Medical University (No. D2016007), The Natural Science Foundation for the Jiangsu Higher Education Institutions of China (No. 16KJB180028), Innovative and Entrepreneurial Talent Scheme of Jiangsu Province (2017), The Natural Science Foundation of the Jiangsu Higher Education Institutions of China (No. 17KJB360014).

## Author Contribution statement

B.G. and Z.Z. designed the study; Z.Z. F.G. and L.W. carried out the experimental work; L.W. Y.L and J.S. analyzed the data; H.Z. and Y.C. prepared the figures and the supplementary tables; P.M. and B.G. do the language modification. Z.Z. and L.W. wrote the manuscript. All authors read and approved the final manuscript.

## Supplementary information

### Competing Interests statement

The authors declare no competing interests.

**Supplementary Figure 1.**
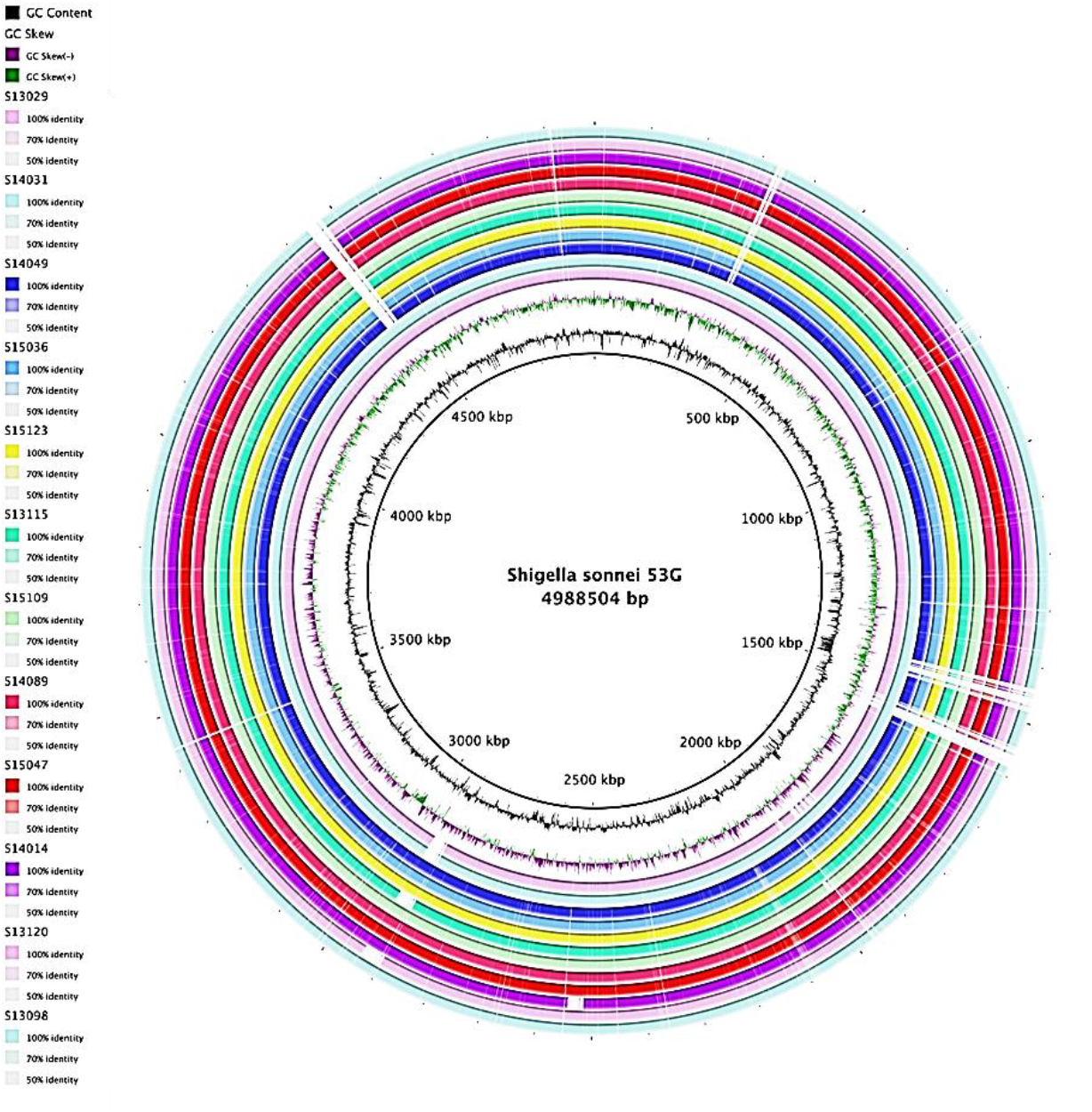
Genome comparison of 12 isolated *S. sonnei* strains against reference genome *S. sonnei* 53G generated by BRIG 0.95. The inner cycle (black) represents the complete genome of the reference strain and the shade of each colors denote the similarities between each strain with reference strain. GC content and GC skew (+/-) were illustrated in-between.

**Supplementary Figure 2.**
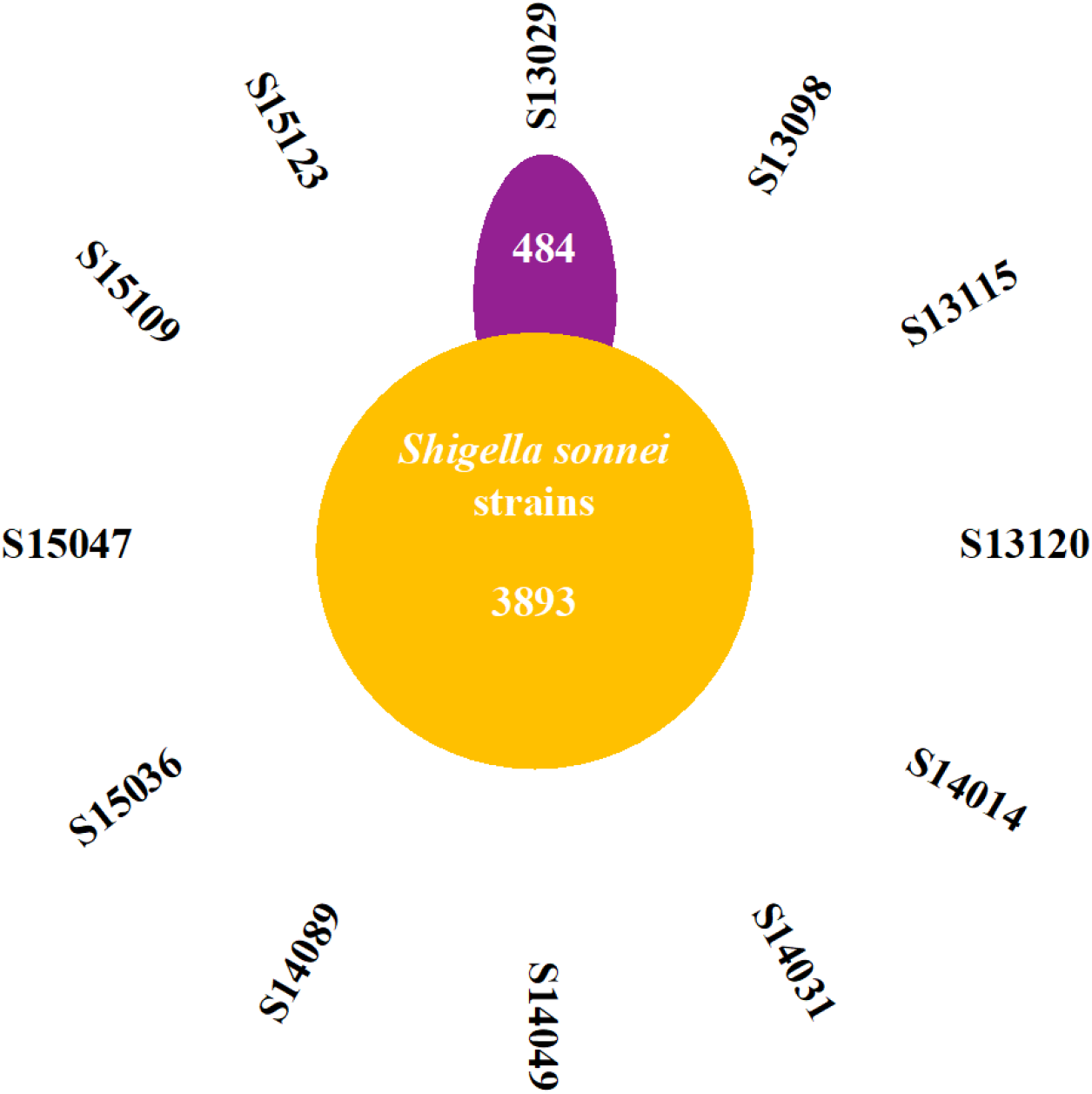
Pangenome analysis of 12 clinically isolated *Shigella sonnei* strains with different antibiotic resistant profiles. A total of 3893 core genes (99% <= strains <= 100%) were shared by all strains while there are 5608 total genes(0% <= strains <= 100%) in all the strains. S13029 and S14031 are sensitive strains while other ten strains have multiple antibiotic resistance to more than 5 antibiotics.

**Supplementary Figure 3.**
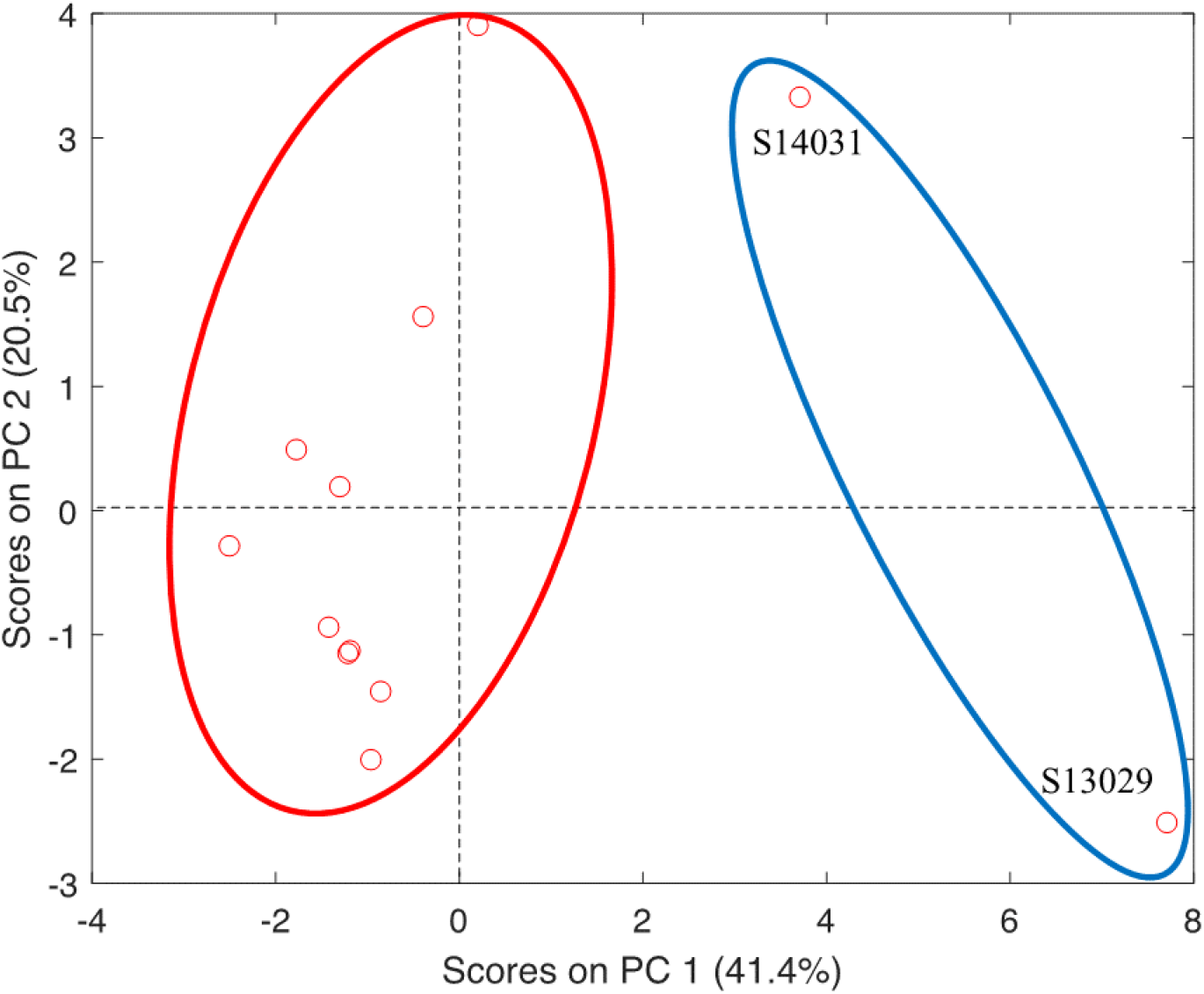
Principal component analysis (PCA) of the relationship between antibiotic resistance profiles and virulence factors in 12 sequenced Shigella sonnei strains. S13029 and S14031 are antibiotics-sensitive strains while the rest strains are multi-drug resistant. (MDR) The two sensitive strains are apparently isolated from other MDR strains.

**Supplementary Table 1.** Comparison of 12 *S. sonnei* strains based on antibiotic resistance profiles and key genome annotation parameters. The 12 strains were isolated from 4 municipal cities, Nanjing, Suzhou, Wuxi, and Zhenjiang, the economy of which are above the average level of China and are considered the best among Jiangsu province. Antibiotics tested in this study are Amoxicillin/Clavulanic acid (AMC), Ceftiophene (CFT), Cefotaxime (CTX), Gentamicin (GEN), Nalidixic acid (NAL), Norfloxacin (NOR), Tetracycline (TBT), and compound Sulfamethoxazole (SMZ).

| Strain ID | Region | AMC | CFT | CTX | GEN | NAL | NOR | TBT | SMZ | MDR | Contig | Bps | CDS | tmRNA | tRNA | rRNA |
| --- | --- | --- | --- | --- | --- | --- | --- | --- | --- | --- | --- | --- | --- | --- | --- | --- |
| S13029 | Zhenjiang | S | S | S | S | S | S | S | S | 0 | 449 | 4490117 | 4328 | 1 | 83 | 5 |
| S13098 | Zhenjiang | R | R | R | R | R | S | R | R | 8 | 473 | 4642881 | 4505 | 1 | 77 | 4 |
| S13115 | Suzhou | S | R | R | R | R | S | S | R | 6 | 409 | 4530601 | 4403 | 1 | 79 | 5 |
| S13120 | Wuxi | I | R | R | R | R | S | R | R | 7 | 406 | 4534710 | 4401 | 1 | 80 | 5 |
| S14014 | Zhenjiang | R | R | R | S | R | S | R | R | 7 | 409 | 4526456 | 4407 | 1 | 80 | 5 |
| S14031 | Nanjing | S | S | S | S | S | S | S | R | 1 | 427 | 4568178 | 4414 | 1 | 76 | 5 |
| S14049 | Wuxi | S | I | S | R | R | S | R | R | 5 | 477 | 4756429 | 4646 | 1 | 77 | 5 |
| S14089 | Suzhou | S | R | R | R | R | S | R | R | 7 | 472 | 4726659 | 4599 | 1 | 80 | 5 |
| S15036 | Nanjing | S | R | R | S | R | S | R | R | 6 | 429 | 4553996 | 4425 | 1 | 78 | 5 |
| S15047 | Zhenjiang | S | R | R | R | R | S | R | R | 7 | 420 | 4606954 | 4484 | 1 | 78 | 5 |
| S15109 | Suzhou | S | R | R | R | R | S | R | R | 7 | 587 | 4541154 | 4378 | 1 | 80 | 5 |
| S15123 | Wuxi | S | R | R | S | R | S | R | R | 6 | 425 | 4645117 | 4528 | 1 | 80 | 5 |

\*MDR: the total number of multi-drug resistance; Bps: base pairs; CDS: coding sequences; S: sensitive (marked red); R: resistant; I: intermittent.

**Supplementary Table 2**: Comparing sensitive strains with resistant strains, unique genes associated with the two groups were identified and annotated based on sequence homology. (in a separate document)

**Supplementary Table 3.** Unique genes specifically associated with *Shigella sonnei* sensitive strains (26 genes) and resistant strains (39 genes). Genes with no assigned specific names are not included in the list. Functions are assigned based on sequence homology.

| Sensitive Strains | Function |
| --- | --- |
| bdm | Biofilm-dependent modulation protein |
| betA | Oxygen-dependent choline dehydrogenase |
| betB_1 | NAD/NADP-dependent betaine aldehyde dehydrogenase |
| betI | HTH-type transcriptional regulator |
| betT | Choline transport protein |
| elaD_1 | Deubiquitinase |
| fimD_1 | Outer membrane usher FimD-like protein |
| fimD_2 | Outer membrane usher FimD-like protein |
| fimD_3 | Outer membrane usher FimD-like protein |
| fimD_4 | Outer membrane usher FimD-like protein |
| fimF | Fimbrial protein |
| fimG | Fimbrial morphology protein |
| fimH_1 | Adhesin |
| flhE | Flagellar biosynthesis protein |
| focC_2 | Chaperone protein |
| gpFI_1 | Putative prophage major tail sheath protein |
| intS_4 | Integrase |
| ompX_2 | Outer membrane protein X |
| pdeL | Cyclic di-GMP phosphodiesterase |
| sat | Serine protease sat autotransporter |
| uacT_1 | Uric acid transporter |
| yahB_2 | Putative HTH-type transcriptional regulator |
| ybdO_1 | Putative HTH-type transcriptional regulator |
| ydiO | Putative fimbrial chaperone |

| ynfE | Putative dimethyl sulfoxide reductase chain |
| --- | --- |
| yraI | Putative fimbrial chaperone |
| Resistant Strains | Function |
| ail | Attachment invasion locus protein |
| ant1 | Streptomycin 3'-adenylyltransferase |
| citC_2 | [Citrate [pro-3S]-lyase] ligase |
| clpP_2 | ATP-dependent Clp protease proteolytic subunit |
| dhfrI | Dihydrofolate reductase type 1 |
| dmlR_2 | HTH-type transcriptional regulator |
| elfC_4 | Putative outer membrane usher protein |
| erfK_1 | Putative L,D-transpeptidase |
| eutB_1 | Ethanolamine ammonia-lyase heavy chain |
| fhuE_1 | FhuE receptor |
| fucI_2 | L-fucose isomerase |
| gabP | GABA permease |
| gatA_1 | PTS system galactitol-specific EIIA component |
| gspA_1 | Putative general secretion pathway protein A |
| gspB | Putative general secretion pathway protein B |
| gspE | Putative type II secretion system protein E |
| hcaR_2 | Hca operon transcriptional activator |
| hyuA_1 | D-phenylhydantoinase |
| mhpC_2 | 2-hydroxy-6-oxononadienedioate/2-hydroxy-6-oxononatrienedioate hydrolase |
| mngB_2 | Mannosylglycerate hydrolase |
| murP_2 | PTS system N-acetylmuramic acid-specific EIIBC component |
| puuC_2 | NADP/NAD-dependent aldehyde dehydrogenase |
| rhaR_2 | HTH-type transcriptional activator |
| tnsA | Transposon Tn7 transposition protein |
| tnsB | Transposon Tn7 transposition protein |
| tnsC | Transposon Tn7 transposition protein |
| tnsE | Transposon Tn7 transposition protein |
| torZ_2 | Trimethylamine-N-oxide reductase 2 |
| umuC_1 | Protein UmuC |
| umuD_1 | Protein UmuD |
| wecD_2 | dTDP-fucosamine acetyltransferase |
| xerC_2 | Tyrosine recombinase |
| xerD_2 | Tyrosine recombinase |
| yadC_3 | Putative fimbrial-like protein |
| ycaM_2 | Inner membrane transporter |
| ycjP_2 | Inner membrane ABC transporter permease protein |
| ygeA | Putative racemase |
| ygfK_2 | Putative oxidoreductase |
| ypdA_2 | Sensor histidine kinase |

**Supplementary Table 4:** The detailed description of assembly results. (in a separate document)

