## Supplementary Table 4 for "Comparative genome analysis of 12 *Shigella sonnei* strains: virulence, resistance, and their interactions"

### Summary of Assembly

#### Sample: S13029

Before assembling, we used k-mer analysis to estimate the size of genome (the assemble result was the real genome size), the degree of heterozygosis and the degree of duplication. The result exhibited that the genome size of sample S13029 was 4.76 Mb. The detail information was shown in figure below.

Based on the assemble result of sample S13029, we found that the genome size was 4,490,790 bp, GC content was 50.69%, the number of scaffold was 419, and the number of contig was 449.

**Table1 Assembling statistics of sample S13029**

| | Scaffold | Contig |
|---|---|---|
| **Total Number (#)** | 419 | 449 |
| **Total Length (bp)** | 4,490,790 | 4,490,117 |
| **N50 (bp)** | 23,418 | 21,183 |
| **N90 (bp)** | 5,565 | 5,080 |
| **Max Length (bp)** | 88,256 | 88,256 |
| **Min Length (bp)** | 512 | 270 |
| **GC Content (%)** | 50.69 | 50.69 |

Note: The second column is the statistic of scaffold longer than 500 bp, and the third column is the statistic of contig got by breaking scaffold (>=500 bp) from the second column with N.

#### Sample: S13034

Before assembling, we used k-mer analysis to estimate the size of genome (the assemble result was the real genome size), the degree of heterozygosis and the degree of duplication. The result exhibited that the genome size of sample S13034 was 4.91 Mb. The detail information was shown in figure below.

Based on the assemble result of sample S13034, we found that the genome size was 4,577,846 bp, GC content was 50.88%, the number of scaffold was 415, and the number of contig was 426.

**Table 2 Assembling statistics of sample S13034**

| | Scaffold | Contig |
|---|---|---|
| **Total Number (#)** | 415 | 426 |
| **Total Length (bp)** | 4,577,846 | 4,577,344 |
| **N50 (bp)** | 25,284 | 24,131 |

| | | |
|---|---|---|
| **N90 (bp)** | 5,461 | 5,221 |
| **Max Length (bp)** | 88,259 | 88,259 |
| **Min Length (bp)** | 511 | 222 |
| **GC Content (%)** | 50.88 | 50.88 |

Note: The second column is the statistic of scaffold longer than 500 bp, and the third column is the statistic of contig got by breaking scaffold (>=500 bp) from the second column with N.

**Sample: S13098**

Before assembling, we used k-mer analysis to estimate the size of genome (the assemble result was the real genome size), the degree of heterozygosis and the degree of duplication. The result exhibited that the genome size of sample S13098 was 5.15 Mb. The detail information was shown in figure below.

Based on the assemble result of sample S13098, we found that the genome size was 4,643,542 bp, GC content was 50.91%, the number of scaffold was 433, and the number of contig was 473.

**Table 3 Assembling statistics of sample S13098**

| | **Scaffold** | **Contig** |
|---|---|---|
| **Total Number (#)** | 433 | 473 |
| **Total Length (bp)** | 4,643,542 | 4,642,881 |
| **N50 (bp)** | 25,309 | 23,337 |
| **N90 (bp)** | 5,062 | 4,661 |
| **Max Length (bp)** | 88,259 | 88,259 |
| **Min Length (bp)** | 513 | 241 |
| **GC Content (%)** | 50.91 | 50.91 |

Note: The second column is the statistic of scaffold longer than 500 bp, and the third column is the statistic of contig got by breaking scaffold (>=500 bp) from the second column with N.

**Sample: S13109**

Before assembling, we used k-mer analysis to estimate the size of genome (the assemble result was the real genome size), the degree of heterozygosis and the degree of duplication. The result exhibited that the genome size of sample S13109 was 4.07 Mb. The detail information was shown in figure below.

Based on the assemble result of sample S13109, we found that the genome size was 4,388,287 bp, GC content was 50.66%, the number of scaffold was 419, and the number of contig was 472.

**Table 4 Assembling statistics of sample S13109**

| | Scaffold | Contig |
|---|---|---|
| Total Number (#) | 419 | 472 |
| Total Length (bp) | 4,388,287 | 4,387,425 |
| N50 (bp) | 24,361 | 22,591 |
| N90 (bp) | 5,244 | 4,730 |
| Max Length (bp) | 95,279 | 95,279 |
| Min Length (bp) | 505 | 205 |
| GC Content (%) | 50.66 | 50.66 |

Note: The second column is the statistic of scaffold longer than 500 bp, and the third column is the statistic of contig got by breaking scaffold (>=500 bp) from the second column with N.

**Sample: S13115**

Before assembling, we used k-mer analysis to estimate the size of genome (the assemble result was the real genome size), the degree of heterozygosis and the degree of duplication. The result exhibited that the genome size of sample S13115 was 4.89 Mb. The detail information was shown in figure below.

Based on the assemble result of sample S13115, we found that the genome size was 4,531,180 bp, GC content was 50.84%, the number of scaffold was 394, and the number of contig was 409.

**Table 5 Assembling statistics of sample S13115**

| | Scaffold | Contig |
|---|---|---|
| Total Number (#) | 394 | 409 |
| Total Length (bp) | 4,531,180 | 4,530,601 |
| N50 (bp) | 25,309 | 25,284 |
| N90 (bp) | 5,794 | 5,419 |
| Max Length (bp) | 88,259 | 88,259 |
| Min Length (bp) | 511 | 222 |
| GC Content (%) | 50.84 | 50.84 |

Note: The second column is the statistic of scaffold longer than 500 bp, and the third column is the statistic of contig got by breaking scaffold (>=500 bp) from the second column with N.

**Sample: S13120**

Before assembling, we used k-mer analysis to estimate the size of genome (the assemble result was the real genome size), the degree of heterozygosis and the degree of duplication. The result exhibited that the genome size of sample S13120 was 4.89 Mb. The detail information was shown in figure below.

Based on the assemble result of sample S13120, we found that the genome size was 4,535,087 bp, GC content was 50.76%, the number of scaffold was 398, and the number of contig was 406.

**Table 6 Assembling statistics of sample S13120**

| | Scaffold | Contig |
|---|---|---|
| **Total Number (#)** | 398 | 406 |
| **Total Length (bp)** | 4,535,087 | 4,534,710 |
| **N50 (bp)** | 25,309 | 25,284 |
| **N90 (bp)** | 5,795 | 5,795 |
| **Max Length (bp)** | 88,259 | 88,259 |
| **Min Length (bp)** | 512 | 222 |
| **GC Content (%)** | 50.76 | 50.76 |

Note: The second column is the statistic of scaffold longer than 500 bp, and the third column is the statistic of contig got by breaking scaffold (>=500 bp) from the second column with N.

**Sample: S14014**

Before assembling, we used k-mer analysis to estimate the size of genome (the assemble result was the real genome size), the degree of heterozygosis and the degree of duplication. The result exhibited that the genome size of sample S14014 was 4.76 Mb. The detail information was shown in figure below.

Based on the assemble result of sample S14014, we found that the genome size was 4,526,793 bp, GC content was 50.77%, the number of scaffold was 395, and the number of contig was 409.

**Table 7 Assembling statistics of sample S14014**

| | Scaffold | Contig |
|---|---|---|
| **Total Number (#)** | 395 | 409 |

| | | |
|---|---|---|
| **Total Length (bp)** | 4,526,793 | 4,526,456 |
| **N50 (bp)** | 25,309 | 25,018 |
| **N90 (bp)** | 5,793 | 5,461 |
| **Max Length (bp)** | 88,259 | 88,259 |
| **Min Length (bp)** | 511 | 264 |
| **GC Content (%)** | 50.77 | 50.77 |

Note: The second column is the statistic of scaffold longer than 500 bp, and the third column is the statistic of contig got by breaking scaffold (>=500 bp) from the second column with N.

**Sample: S14031**

Before assembling, we used k-mer analysis to estimate the size of genome (the assemble result was the real genome size), the degree of heterozygosis and the degree of duplication. The result exhibited that the genome size of sample S14031 was 4.76 Mb. The detail information was shown in figure below.

Based on the assemble result of sample S14031, we found that the genome size was 4,568,747 bp, GC content was 50.69%, the number of scaffold was 397, and the number of contig was 427.

**Table 8 Assembling statistics of sample S14031**

| | **Scaffold** | **Contig** |
|---|---|---|
| **Total Number (#)** | 397 | 427 |
| **Total Length (bp)** | 4,568,747 | 4,568,178 |
| **N50 (bp)** | 27,037 | 25,717 |
| **N90 (bp)** | 6,081 | 5,794 |
| **Max Length (bp)** | 88,235 | 81,068 |
| **Min Length (bp)** | 500 | 211 |
| **GC Content (%)** | 50.69 | 50.69 |

Note: The second column is the statistic of scaffold longer than 500 bp, and the third column is the statistic of contig got by breaking scaffold (>=500 bp) from the second column with N.

After genome assembling, the GC distribution of sample S14031 was obtained by GC-Depth analysis. The result was shown below.

**Sample: S14049**

Before assembling, we used k-mer analysis to estimate the size of genome (the assemble result was the real genome size), the degree of heterozygosis and the degree of duplication. The result exhibited that the genome size of sample S14049 was 5.28 Mb. The detail information was shown in figure below.

Based on the assemble result of sample S14049, we found that the genome size was 4,757,139 bp, GC content was 50.56%, the number of scaffold was 459, and the number of contig was 477.

**Table 9 Assembling statistics of sample S14049**

| | Scaffold | Contig |
|---|---|---|
| **Total Number (#)** | 459 | 477 |
| **Total Length (bp)** | 4,757,139 | 4,756,429 |
| **N50 (bp)** | 25,284 | 24,131 |
| **N90 (bp)** | 4,589 | 4,513 |
| **Max Length (bp)** | 88,259 | 88,259 |
| **Min Length (bp)** | 504 | 222 |
| **GC Content (%)** | 50.56 | 50.56 |

Note: The second column is the statistic of scaffold longer than 500 bp, and the third column is the statistic of contig got by breaking scaffold (>=500 bp) from the second column with N.

**Sample: S14089**

Before assembling, we used k-mer analysis to estimate the size of genome (the assemble result was the real genome size), the degree of heterozygosis and the degree of duplication. The result exhibited that the genome size of sample S14089 was 5.28 Mb. The detail information was shown in figure below.

Based on the assemble result of sample S14089, we found that the genome size was 4,728,784 bp, GC content was 50.6%, the number of scaffold was 433, and the number of contig was 472.

**Table 10 Assembling statistics of sample S14089**

| | Scaffold | Contig |
|---|---|---|
| **Total Number (#)** | 433 | 472 |
| **Total Length (bp)** | 4,728,784 | 4,726,659 |
| **N50 (bp)** | 25,609 | 25,017 |
| **N90 (bp)** | 4,991 | 4,589 |
| **Max Length (bp)** | 88,259 | 88,259 |

| | | |
|---|---|---|
| **Min Length (bp)** | 511 | 206 |
| **GC Content (%)** | 50.6 | 50.6 |

Note: The second column is the statistic of scaffold longer than 500 bp, and the third column is the statistic of contig got by breaking scaffold (>=500 bp) from the second column with N.

**Sample: S15036**

Before assembling, we used k-mer analysis to estimate the size of genome (the assemble result was the real genome size), the degree of heterozygosis and the degree of duplication. The result exhibited that the genome size of sample S15036 was 5.28 Mb. The detail information was shown in figure below.

Based on the assemble result of sample S15036, we found that the genome size was 4,555,699 bp, GC content was 50.78%, the number of scaffold was 398, and the number of contig was 429.

**Table 11 Assembling statistics of sample S15036**

| | **Scaffold** | **Contig** |
|---|---|---|
| **Total Number (#)** | 398 | 429 |
| **Total Length (bp)** | 4,555,699 | 4,553,996 |
| **N50 (bp)** | 25,309 | 23,764 |
| **N90 (bp)** | 5,795 | 5,461 |
| **Max Length (bp)** | 88,259 | 88,259 |
| **Min Length (bp)** | 511 | 222 |
| **GC Content (%)** | 50.78 | 50.78 |

Note: The second column is the statistic of scaffold longer than 500 bp, and the third column is the statistic of contig got by breaking scaffold (>=500 bp) from the second column with N.

**Sample: S15047**

Before assembling, we used k-mer analysis to estimate the size of genome (the assemble result was the real genome size), the degree of heterozygosis and the degree of duplication. The result exhibited that the genome size of sample S15047 was 5.15 Mb. The detail information was shown in figure below.

Based on the assemble result of sample S15047, we found that the genome size was 4,607,635 bp, GC content was 50.85%, the number of scaffold was 400, and the number of contig was 420.

**Table 12 Assembling statistics of sample S15047**

| | Scaffold | Contig |
|---|---|---|
| **Total Number (#)** | 400 | 420 |
| **Total Length (bp)** | 4,607,635 | 4,606,954 |
| **N50 (bp)** | 25,674 | 25,284 |
| **N90 (bp)** | 5,793 | 5,225 |
| **Max Length (bp)** | 88,259 | 88,259 |
| **Min Length (bp)** | 511 | 222 |
| **GC Content (%)** | 50.85 | 50.85 |

Note: The second column is the statistic of scaffold longer than 500 bp, and the third column is the statistic of contig got by breaking scaffold (>=500 bp) from the second column with N.

**Sample: S15123**

Before assembling, we used k-mer analysis to estimate the size of genome (the assemble result was the real genome size), the degree of heterozygosis and the degree of duplication. The result exhibited that the genome size of sample S15123 was 5.15 Mb. The detail information was shown in figure below.

Based on the assemble result of sample S15123, we found that the genome size was 4,645,679 bp, GC content was 50.69%, the number of scaffold was 401, and the number of contig was 425.

**Table 13 Assembling statistics of sample S15123**

| | Scaffold | Contig |
|---|---|---|
| **Total Number (#)** | 401 | 425 |
| **Total Length (bp)** | 4,645,679 | 4,645,117 |
| **N50 (bp)** | 25,674 | 25,284 |
| **N90 (bp)** | 5,793 | 5,293 |
| **Max Length (bp)** | 88,259 | 88,259 |
| **Min Length (bp)** | 511 | 221 |
| **GC Content (%)** | 50.69 | 50.69 |

Note: The second column is the statistic of scaffold longer than 500 bp, and the third column is the statistic of contig got by breaking scaffold (>=500 bp) from the second column with N.
